# Enhanced Irrigation during Extreme Heat Events Preserves Anthocyanins in Cabernet Sauvignon

**DOI:** 10.64898/2026.05.26.727714

**Authors:** James R. Campbell, Martina Galeano, Andrew J. McElrone, Luis Sanchez, Nick Dokoozlian, Sophia Bagshaw, Andrew L. Waterhouse, Elisabeth J. Forrestel

## Abstract

Globally, heat waves (HWs) have become more frequent, intense, and prolonged, with extreme temperatures that reduce quality or result in crop loss in wine grapes. Irrigation prior to or during HWs is one of the most common means of mitigating damage to vines and berries. However, the effect of this practice on flavonoids is not well established. Red wine quality is directly impacted by phenolics, including anthocyanins and proanthocyanidins (PAs). This study was conducted over three vintages (2019-2021) in a commercial vineyard to evaluate the impact of supplemental irrigation – applied before and during HWs – on winegrape chemistry in Cabernet Sauvignon. Results demonstrated that supplemental irrigation significantly reduced anthocyanin loss compared to a control treatment maintained at 60% evapotranspiration (ET), and that pre-HW irrigation can mitigate some of the deleterious effects on classes of flavonoids important to red wine quality. Furthermore, applying excessive water (3x, or 180% ET) had no additional beneficial effects on flavonoids relative to a moderate supplemental application (2x, or 120% ET).

## Introduction

Heat waves (HWs) are increasing in intensity, frequency, and duration, significantly impacting agricultural production and fruit quality.^1,2^ Combined with growing water scarcity, HWs pose substantial challenges for agriculture, making it critical to understand how water conservation during extreme heat events can be balanced with maintaining fruit quality. Regulated deficit irrigation (RDI), which induces mild water stress, is commonly used to reduce canopy vigor and yield, creating a more favorable microclimate for grape clusters, smaller berries, and improved fruit composition.^3–5^ Standard RDI in the grape and wine industry is typically applied at 60–70% of evapotranspiration (ET), allowing fruit to reach full sugar maturity while promoting anthocyanin synthesis during ripening in red winegrapes.^6^

Elevated temperatures significantly affect the production of primary and secondary berry metabolites through both whole-vine physiological responses and direct fruit exposure.^7^. Primary metabolites relevant to wine production include total soluble solids (Brix), titratable acidity (TA), and pH. Warmer growing conditions accelerate berry development, increasing rates of sugar loading and ultimately sugar concentration — and therefore alcohol levels in wine.^7–9^ However, extreme heat can delay sugar accumulation by reducing carbon assimilation, thus slowing ripening.^10^

RDI has also been associated with decreased TA, attributed to greater malic acid consumption.^4,5^ By reducing canopy vigor, RDI increases solar radiation exposure and canopy temperatures, enhancing respiration. Vine water status is closely tied to sugar accumulation: RDI can promote sugar translocation to berries, while severe deficit inhibits photosynthesis and carbon assimilation, ultimately reducing fruit sugar content.^11^ Importantly, the beneficial effects of RDI on berry quality may not hold under future climate conditions, as combined heat and water stress are likely to interact in ways detrimental to berry and wine quality.^12^

Acidity – measured as pH and TA – is a key indicator of fruit quality. (L)-Tartaric and (L)-malic acids are synthesized in green berries and stored in mesocarp and skin cells.^13^ Pre-veraison, warmer conditions favor (L)-malic acid synthesis, while post-veraison, elevated temperatures accelerate (L)-malate breakdown, lowering TA and raising must pH.^12^ At the onset of ripening, grapevines shift from sugars to (L)-malate as the primary respiratory substrate; this breakdown increases at higher temperatures.^4,14^ (L)-Tartrate content, by contrast, remains stable — observed concentration decreases are attributed to dilution from berry growth and expansion.^13^ Temperature effects on pH and TA appear to be cultivar-dependent, and further research on the dynamics of (L)-malate, (L)-tartrate, potassium, and sodium is needed to fully understand environmental and genetic influences on these properties.^15–17^

RDI influences secondary berry chemistry through several mechanisms. First, reduced berry size increases the skin-to-pulp ratio, concentrating phenolic compounds. Second, RDI upregulates genes in the phenylpropanoid pathway, enhancing anthocyanin synthesis.^11,18,19^ The effect of RDI on skin proanthocyanidins (PAs), or condensed tannins, appears to be driven primarily by berry size reduction rather than changes in biosynthesis.^6,11^ Pre-veraison RDI has been shown to not affect PA content directly,^20^ though indirect effects on canopy size are impactful, as increased light exposure on fruit pre-veraison can stimulate PA synthesis.^21,22^

Among secondary metabolites, flavonoids and aromatic compounds are of particular interest for grape and wine quality. The phenylpropanoid pathway governs the biosynthesis of numerous grape phenolics.^23–25^ Flavonoids share a characteristic C6–C3–C6 structure and are ubiquitous in the plant kingdom, playing especially important roles in red winegrape varieties. They contribute to a range of chemical and sensory properties in wine, including color, bitterness, astringency, and aging potential. The major flavonoid subclasses in grape skins — anthocyanins, flavonols, flavan-3-ols, and proanthocyanidins — are each variably affected by elevated temperatures. ^26^

Anthocyanins are the pigment molecules responsible for red wine color. Their synthesis begins at veraison, peaks near maturity, and may subsequently decline ^27,28^. Anthocyanin accumulation appears to be driven primarily by temperature rather than light, provided photon flux exceeds 100 μmol m ² s ¹.^29–31^ Earlier work suggested optimal synthesis near 30°C with inhibition beginning around 35°C,^32^ while more recent growth chamber studies show inhibition can occur at 30°C.^33^ Both studies identified temperature-driven divergence in transcription factor activity leading to reduced anthocyanin synthesis. Beyond inhibited synthesis, elevated temperatures also appear to promote anthocyanin degradation via peroxidases.^27,34^ In Cabernet Sauvignon, Mori et al. found that high temperatures did not inhibit biosynthesis per se, but that anthocyanin loss was primarily attributable to chemical and enzymatic degradation.^35^ Sandras and Moran further proposed a thermal decoupling between total soluble solids (TSS) and anthocyanins, demonstrating that vines grown under warmer conditions and varying irrigation regimes contained lower anthocyanin levels at equivalent ripeness compared to controls.^36^

Flavonols are a class of phenolics primarily associated with light exposure and are well documented to accumulate in sun-exposed fruit. ^37^ Functioning as a “sunscreen” and antioxidant, they help protect tissue from oxidative damage in high-light environments. They also contribute to the color of young wines as copigments, through intermolecular non-covalent interactions.^38^ Flavan-3-ols are the most abundant flavonoids in grapes and wines; more than 50% exist as oligomers and polymers (PAs) formed through flavan-3-ol condensation. ^39,40^ Monomeric flavan-3-ols contribute to bitterness, while polymeric PAs are associated with astringency. These compounds also react with anthocyanins to form stable pigments and have notable antioxidant properties.^27,41,42^

PAs are found in both skin and seeds, located in cell vacuoles or bound to cell wall material (CWM).^28,43^ Synthesis begins at flowering, continues through early berry development, and peaks around veraison.^40,44^ Post-veraison, PA concentrations decline throughout ripening — likely due to reduced extractability from CWM interactions rather than degradation.^45,46^ The effects of elevated temperatures on skin and seed PAs have produced inconsistent findings. Cohen et al. observed a linear relationship between heat accumulation during early berry development and skin PA content, but found no such relationship under diurnal temperature compression.^47,48^ Gouot et al. reported an increase in total skin PA concentration in response to short pre-veraison heat stress, but no difference in skin PA content at maturity.^49^ Heat applied post-veraison has generally shown no effect on PA levels, possibly because synthesis has ceased by that point, or because PAs are more thermally stable than other flavonoids due to the higher activation energy required for heterocyclic ring degradation.^27,50^ Field studies examining heat exposure across the entire growing season found decreased total PA content and concentration at maturity, driven primarily by seed PA reductions, while skin PAs were largely unaffected. ^12^.

Collectively, prior work indicates that elevated temperatures and variable irrigation regimes under controlled or semi-controlled conditions can reduce flavonoid content in winegrapes, particularly anthocyanins, which are especially susceptible to thermal degradation. ^34,35,49^ While there have been studies of utilizing variable misting systems and sustained irrigation regimes (e.g.,^51,52^), no field experiments have directly evaluated whether targeted increases in irrigation — a primary heat mitigation strategy already widely employed by growers — can protect berry flavonoid composition during naturally occurring heat extremes. We hypothesized that increasing water availability before and during a heat wave would mitigate flavonoid, and more specifically anthocyanin loss, in red winegrapes (*V. vinifera* subsp. *vinifera* cv. Cabernet Sauvignon), thereby limiting color loss and potential impacts on wine quality during the ripening phase.

## Materials & Methods

### Field site

This study was conducted across the 2019, 2020, and 2021 growing seasons in a commercial vineyard in Herald, CA, USA. The site, located in California’s San Joaquin Valley, consisted of a 23-acre block of 9-year-old *Vitis vinifera* cv. Cabernet Sauvignon grafted onto 1103P rootstock (Figure S1). Rows were oriented east–west with vine spacing of 1.8 m and row spacing of 3 m, trained to a bilateral cordon with a single high wire. The vineyard block was divided into 98 subplots of 30 m × 30 m, each served by a variable rate drip irrigation (VRDI) system that allowed differential irrigation to be applied to individual subplots. Irrigation was scheduled throughout the growing season by calculating crop evapotranspiration (ETc) from reference evapotranspiration (ETo) obtained from a nearby weather station combined with a dynamic, modified crop coefficient (Kc) approach. Kc values were derived from remotely sensed normalized difference vegetation index (NDVI) data specific to each 30 × 30 m zone, along with empirical parameters representing the expected water demand of a non-stressed vineyard .^53^

### Defining heat waves and irrigation application

Based on historical weather data, an extreme heat event (heat wave; HW) was defined as three or more consecutive days with a daily maximum temperature exceeding 38°C. This threshold was determined using 1980–2010 climate data as the baseline, specifically the 95th percentile of maximum temperatures during the growing season (April–October), consistent with the principle that HWs are defined relative to local historical conditions. No artificial heating system was used; the study relied entirely on naturally occurring HWs. During the 2019 growing season, two HWs occurred: one in late July and one in mid-August. In 2020, four HWs were recorded: one in late May, one in mid-July, one in mid-August, and one in early September. In 2021, three HWs occurred: one in mid-June, one in early July, and one in early September. Maximum temperatures were recorded by the vineyard weather station (38°17′24″N, 121°7′12″W) associated with the GrapeX project ^54^, located 0.8 km from the field experiment.

In 2019, three subplots (30 m × 30 m) per treatment were randomly selected and distributed across the site using the VRDI system to deliver differential irrigation. In 2020 and 2021, each replicate was expanded to four subplots, forming 60 m × 60 m plots (Figure S1). Evapotranspiration was estimated throughout the season using Landsat-derived NDVI and a crop coefficient approach. ^53^

Differential irrigation treatments were applied only during HWs, beginning one to two days before each event and continuing through the final day. Three treatments were applied: (1) a control maintained at deficit irrigation (60% ETc, no additional water); (2) double the control rate (120% ETc); and (3) triple the control rate (180% ETc). These treatments were used in 2019 and 2020. Outside of HW periods, all treatments were maintained at 60% ETc (Figure 1). In 2021, treatment levels were adjusted to 60%, 90%, and 120% ETc during HWs, based on findings from the previous two seasons and consultation with growers.

**Figure 1.**
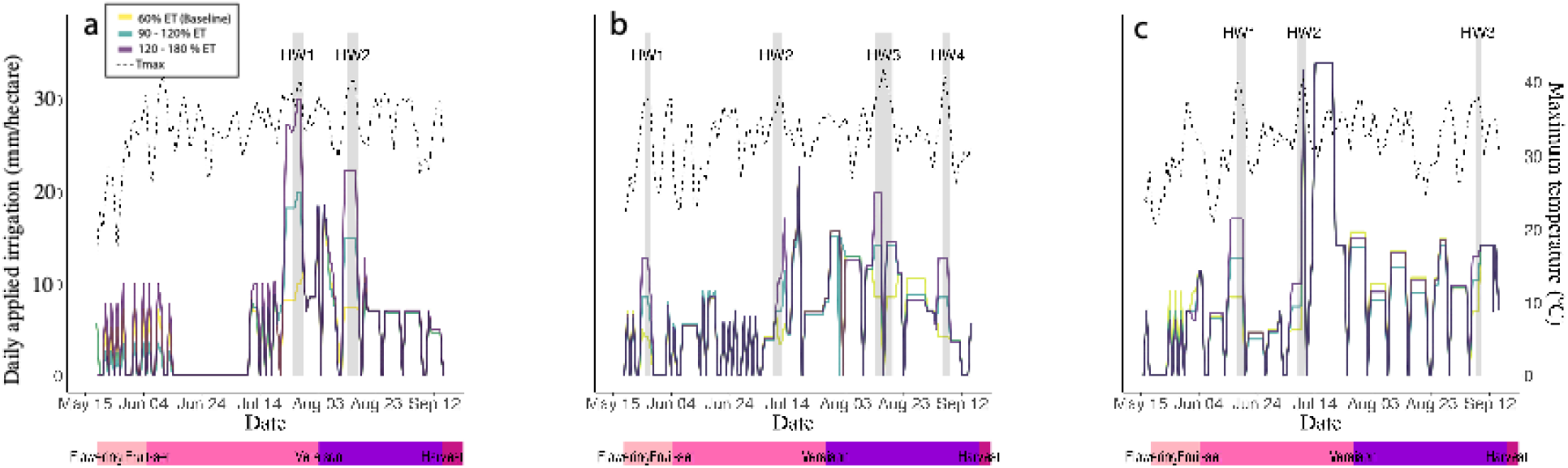
Daily applied irrigation in gallons/acre across each growing season monitored; a) 2019, b) 2020, c) 2021; secondary y-axis represents maximum daily temperature across the growing seasons. The colored lines under each plot illustrate the timing of flowering, fruit set, veraison and harvest.

Based on historical weather data, an extreme heat event was defined as a minimum of three consecutive days with the daily maximum temperature exceeding 38° C. Recognizing that HWs are defined relative to local historical conditions, the daily maximum temperature cutoff of 38° C was determined using the 1980-2010 climate as the baseline, the 95% percentile of maximum temperatures during the growing season (April - October) were used to determine temperature above which was considered the beginning of a HW.

No heating system was applied to simulate artificial HWs, rather the study relied on the occurrence of natural HWs. During the 2019 growing season, two HWs took place, one at the end of July and the second one during mid-August, while the 2020 season had a total of four HWs, one at the end of May, one in mid-July, one in mid-August, and one in the beginning of September. As for the 2021 season, three HWs were recorded, one in mid-June, one in early July and the last one at the beginning of September. Maximum temperatures were recorded using the vineyard (38° 17’ 24“ N 121° 7’ 12” W) weather station from the GrapeX project ^54^, which is located 0.8 kilometers from the field experiment. In 2019, three subplots (30 m x 30 m) per treatment were randomly selected and distributed along the site using the VRDI system to provide differential irrigation to the individual plots. In 2020 and 2021, each replicate was expanded to four subplots, or a 60-meter x 60-meter plot (**Figure S1**). Evapotranspiration was estimated throughout the season using Landsat data, normalized difference vegetation index (NDVI) and crop coefficient.^53^

Differential irrigation treatments were applied only when a HW took place and started one or two days before each HW and continued through the last day of the HW. Three irrigation treatments were implemented: 1) a control, or baseline, which was exposed to deficit irrigation and held at 60% ET with no additional water applied, 2) double the baseline (120% ET), and 3) triple the amount of water of the baseline (180% ET) for 2019 and 2020 seasons. Irrigation in all treatments was maintained at a target of 60% ET outside of heatwave events (**Figure 1**). In 2021, irrigation treatments were adjusted to 60% ET, 90% ET, and 120% ET during HWs based on findings from the previous two years of the experiment and consultation with growers.

### Berry Sampling and Chemistry

Berries were sampled at regular intervals from veraison through harvest in 2019 and from fruit set through harvest in 2020 and 2021. Additional sampling was conducted before, during, and after each HW. Nine vines per plot were randomly selected for sampling. From each vine, 40 berries (20 per canopy side) were collected from different clusters and cluster positions, yielding 360 berries per plot per sampling point.

Immediately after sampling, a random subset of 180 berries was divided into three replicates of 60 berries each. Berries were crushed and the resulting juice was centrifuged at 4,200 rpm for 5 minutes. Each juice sample was then analyzed for total soluble solids (TSS) using a Sper Scientific 30051 refractometer (Sper Scientific Ltd., Scottsdale, AZ, USA), pH using an Orion Star A211 pH meter (Thermo Fisher Scientific Inc., Waltham, MA, USA), and titratable acidity (TA) by titration with 0.1 N NaOH (VWR International, Radnor, PA, USA) to an endpoint of pH 8.2.

#### Extraction

Sixty berries were peeled to separate skins from pulp and seeds. Skins were placed in a 100 mL Erlenmeyer flask with 50 mL of 66% (v/v) acetone and homogenized using an Ultra-Turrax T25 (IKA-Werke GmbH, Staufen, Germany) for 1 minute at 13,000 rpm. The homogenate was transferred to a 100 mL opaque jar, and the flask was rinsed with an additional 50 mL of 66% acetone to maximize recovery of skin material. Samples were extracted for 24 h on an orbital shaker at 130 rpm.

Following extraction, samples were transferred to 250 mL centrifuge bottles and centrifuged at 10,000 × g for 10 minutes, then filtered through a 120 mm Büchner funnel with No. 1 Whatman filter paper into a round-bottom flask. Acetone was removed under reduced pressure using a Büchi R-215 rotary evaporator (Büchi Corporation, New Castle, DE, USA) at 80 mbar and 30°C for 10 minutes. The extract was transferred to a graduated cylinder and brought to a final volume of 50 mL with distilled water, then stored at −80°C until analysis.

#### Total Proanthocyanidins Analysis

Proanthocyanidins (PAs) were quantified in triplicate using methyl cellulose precipitation (MCP) as previously described. ^55^ Briefly, 25 µL of extract was added to a 1,200 µL deep-well plate and combined with either 300 µL of 0.04% (w/v) methyl cellulose (Sigma-Aldrich, St. Louis, MO, USA) or 300 µL of water (control). Samples were mixed on a Thermomixer C (Eppendorf AG, Hamburg, Germany) at 1,500 rpm for 5 minutes and allowed to stand for 3 minutes. Then 200 µL of saturated ammonium sulfate was added to prevent re-release of precipitated PA material, followed by water (475 µL for treated samples, 775 µL for controls), and samples were mixed again for 5 minutes. After standing for 10 minutes, the plate was centrifuged at 2,272 × g for 5 minutes (5804R, Eppendorf AG). Aliquots of 200 µL from both treated and control wells were transferred to a 96-well half-area plate (Corning, Corning, NY, USA). PA content was quantified as Δ absorbance at 280 nm (control − treatment) using a linear regression from an (−)-epicatechin standard curve (Sigma-Aldrich). Absorbances were measured simultaneously on a Synergy Neo2 Multi-Mode Reader (BioTek Instruments Inc., Winooski, VT, USA).

#### Anthocyanin and Flavonol Analysis

Anthocyanins and flavonols were analyzed using a previously published HPLC method with modifications ^56^, on an Agilent 1260 system (Santa Clara, CA, USA) equipped with a diode array detector (DAD). Separation was performed on a Phenomenex Kinetex PFP column (150 × 3.0 mm, 2.6 µm particle size) at 50°C with a 10 µL injection volume. Mobile phase A was Milli-Q water with 0.2% TFA; mobile phase B was acetonitrile with 0.2% TFA. The flow rate was 0.5 mL/min. The gradient was as follows: 12.5% B for 0–3 min; linear ramp to 20% B from 3–14 min; 20–27.3% B from 14–26 min; step to 70% B at 26–26.02 min; held at 70% B for 2 minutes; return to 12.5% B at 28–28.02 min; re-equilibration for 4 minutes. Eluting peaks were monitored at 365 nm (flavonols) and 520 nm (anthocyanins). Anthocyanins were quantified against a malvidin-3-*O*-glucoside standard curve (Extrasynthese, Genay, France); flavonols were quantified against a quercetin-3-*O*-glucoside standard curve (Extrasynthese).

## Statistical Analysis

All statistical analyses were performed in R v 3.6.3 using tidyverse, ggplot2, ggpubr, agricolae, factoextra, ggpmisc, and magrittr packages. Code for all analyses is available at https://github.com/mgaleano-ucd/Borden_hills_HWs_complete. One- and two-way ANOVAs with a post-hoc Tukey’s HSD test were used for mean comparisons, with p ≤ 0.05 used as a significance threshold. Error bars and variability shown in tables and graphs are representative of the standard error of mean (SEM).Tables were created in Excel (Microsoft Corporation, Redmond, WA) and figures were in Prism 9 (GraphPad Software Inc., San Diego, CA).

## Results

### Total Soluble Solids, pH, Titratable Acidity, Berry Weights, and Yield

#### TSS

Supplemental irrigation consistently reduced TSS relative to the 60% ET baseline, with treatment separation closely tracking HW occurrence across all years. In 2019 and 2020, TSS differed significantly between the control and supplemental treatments throughout the season (Table S1, S2), though harvest differences were absent in 2019 due to offset picking dates used to match commercial TSS targets (Table 2). In 2020, differences emerged after each HW event and the 60% ET baseline reached its highest relative TSS following HW4, remaining significantly higher at commercial harvest. In 2021, the 60% ET baseline had significantly higher TSS than both supplemental treatments prior to HW2, but no significant differences were observed from August 17th through harvest (Table S3).

#### pH

In all years, the 60% ET baseline trended toward higher pH than the supplemental treatments, with differences emerging and intensifying around HW events. In 2019, pH diverged during HW2 and the 120% ET treatment had significantly lower pH than both other treatments at harvest, while the 180% ET and 60% ET treatments did not differ. In 2020, the 60% ET baseline maintained significantly higher pH from HW3 through commercial harvest (Tables 1 & S2). In 2021, the 90% and 120% ET treatments had significantly lower pH than the baseline from approximately two weeks after HW2 onward, though no significant differences remained at commercial harvest (Tables 2 & S3).

#### TA

TA responses were consistent across years: early in the season the 60% ET baseline tended toward higher TA, but by late season and harvest the supplemental treatments had significantly higher TA than the baseline. In 2019, the 120% ET treatment had significantly higher TA than both other treatments from September 3rd through harvest (Tables 2 & S1). In 2020, the 60% ET baseline had higher TA than the supplemental treatments after HW2, but this was reversed by HW4 and persisted at harvest (Tables 2 & S2). In 2021, TA differences were largely absent until September 8th, when the 60% ET baseline had significantly lower TA; this difference persisted at commercial harvest while pH did not (Tables 2 & S3).

#### Berry weight

Across all three years, supplemental irrigation during HWs consistently produced higher berry weights than the 60% ET baseline, with differences most pronounced at harvest. In 2019, sampling began at veraison so pre-veraison HW effects could not be captured, but both supplemental treatments had significantly higher berry weights than the control from veraison through harvest, with significant differences between the 120% and 180% ET treatments at only two time points (Table 1). In 2020, separation emerged at HW2 and persisted through harvest, with the 120% ET treatment highest following HW3. In 2021, treatments were adjusted to 90% and 120% ET; significant differences between the baseline and supplemental treatments were present before the uniform post-HW irrigation event but largely absent afterward, with only one date with significant differences in weight observed post-veraison (6-Aug).

**Table 1.**
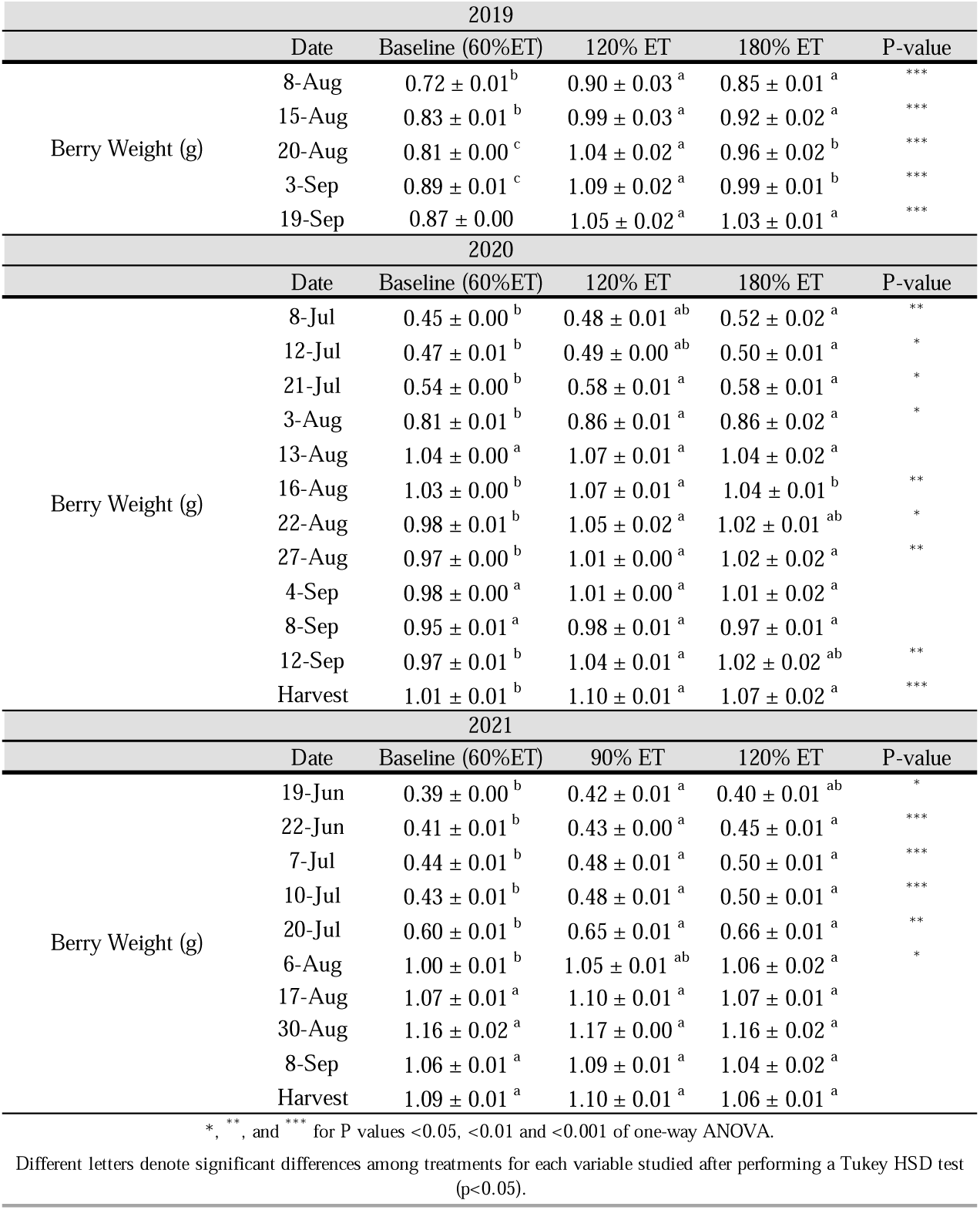
Berry weights measured during the 2019, 2020, and 2021 growing seasons taken from vines exposed to differential irrigation during HWs. Error bars are in SEM. N=9.

#### Yield

Supplemental irrigation increased yield relative to the 60% ET baseline in all three years, though the magnitude and significance varied. In 2019, yield differed significantly between the 60% and 120% ET treatments but not the 60% and 180% ET treatments. In 2020, the baseline had significantly lower yield than both supplemental treatments, with the 120% ET numerically higher than the 180% ET. In 2021, yield differed significantly between the 60% and 90% ET treatments but not between the 60% and 120% ET treatments (Table 2).

**Table 2.** Yield components and primary chemistry of the 2019, 2020, and 2021 growing seasons taken at harvest of vines exposed to differential irrigation prior to and during HWs. Error bars are SEM. N=9.

| 2019 |  |  |  |  |
| --- | --- | --- | --- | --- |
|  | Baseline (60%ET) | 120% ET | 180% ET | P-value |
| **Brix** | **25.62 ± 0.10<sup>a</sup>** | **25.79 ± 0.17 <sup>a</sup>** | **25.51 ± 0.14<sup>a</sup>** |  |
| **pH** | **3.67 ± 0.02<sup>a</sup>** | **3.5 ± 0.03<sup>b</sup>** | **3.66 ± 0.01<sup>a</sup>** | **\*\*\*** |
| **Titrateable acidity (g/L)** | **3.91 ± 0.09<sup>b</sup>** | **4.95 ± 0.16<sup>a</sup>** | **4.27 ± 0.06<sup>b</sup>** | **\*\*\*** |
| **Yield (kg/plant)** | **6.25 ± 0.23<sup>b</sup>** | **8.86 ± 0.63<sup>a</sup>** | **7.7 ± 0.58<sup>ab</sup>** | **\*** |
| 2020 |  |  |  |  |
|  | Baseline (60%ET) | 120% ET | 180% ET | P-value |
| **Brix** | **26.88 ± 0.16<sup>a</sup>** | **25.89 ± 0.23<sup>b</sup>** | **26 ± 0.22<sup>b</sup>** | **\*\*** |
| **pH** | **3.66 ± 0.01<sup>a</sup>** | **3.54 ± 0.02<sup>b</sup>** | **3.54 ± 0.02<sup>b</sup>** | **\*\*\*** |
| **Titrateable acidity (g/L)** | **4.02 ± 0.08<sup>b</sup>** | **4.54 ± 0.13<sup>a</sup>** | **4.86 ± 0.12<sup>a</sup>** | **\*\*\*** |
| **Yield (kg/plant)** | **7.28 ± 0.17<sup>b</sup>** | **8.32 ± 0.2<sup>a</sup>** | **8.24 ± 0.20<sup>a</sup>** | **\*\*\*** |
| 2021 |  |  |  |  |
|  | Baseline (60%ET) | 90% ET | 120% ET | P-value |
| **Brix** | **24.58 ± 0.19<sup>a</sup>** | **24.53 ± 0.18<sup>a</sup>** | **24.66 ± 0.25<sup>a</sup>** |  |
| **pH** | **3.75 ± 0.02<sup>a</sup>** | **3.69 ± 0.02<sup>a</sup>** | **3.72 ± 0.02<sup>a</sup>** |  |
| **Titrateable acidity (g/L)** | **3.20 ± 0.05<sup>b</sup>** | **3.42 ± 0.05<sup>a</sup>** | **3.42 ± 0.04<sup>a</sup>** | **\*\*** |
| **Yield (kg/plant)** | **7.4 ± 0.27<sup>b</sup>** | **8.89 ± 0.29<sup>a</sup>** | **8.33 ± 0.27<sup>ab</sup>** | **\*\*\*** |
| *, **, and *** for P values <0.05, <0.01 and <0.001 of one-way ANOVA. |  |  |  |  |
| Different letters denote significant differences among treatments for each variable studied after performing a Tukey HSD test (p<0.05). |  |  |  |  |

### Monomeric Anthocyanin Content

Across all three years, supplemental irrigation consistently protected anthocyanins from HW-induced degradation, with the 60% ET baseline showing the greatest losses following HW events and the lowest anthocyanin content at harvest. The 120% ET treatment generally performed as well as or better than the highest irrigation treatment, leading to the downward adjustment of treatment levels in 2021.

In 2019, only one post-veraison HW (HW2) met the sampling criteria (Figure 2A). Following HW2, the 60* ET baseline showed a significant decline in total anthocyanins while the 120* ET treatment increased, and the 180* ET plateaued. At harvest, both supplemental treatments had significantly higher anthocyanin content than the 60% ET baseline. In 2020, significant differences emerged following HW3, with the 120% ET recording the highest anthocyanins per berry (Figure 2B). The gap widened after HW4, at which point the 60% ET baseline was significantly lower than both supplemental treatments. Based on these results, the 2021 treatments were adjusted to 90% and 120% ET to optimize water use while maintaining anthocyanin protection. In 2021, the 60% ET baseline had significantly lower anthocyanins than the supplemental treatments as early as the second post-veraison sampling point, persisting through HW3 (Figure 2C). At commercial harvest, no significant differences were observed between the two supplemental treatments.

**Figure 2.**
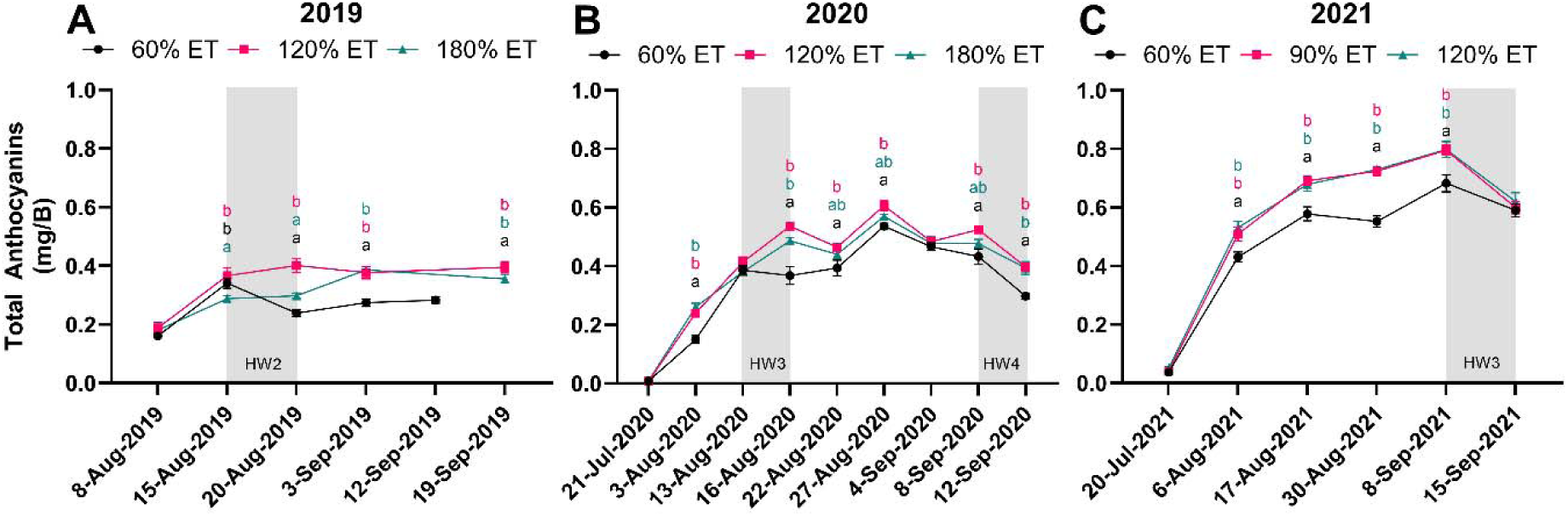
Total anthocyanin content (milligrams of malvidin-3-glucoside equivalents per berry) throughout ripening during the 2019 (A) and 2020 (B) seasons for the 60% ET baseline, 120% ET, and 180% ET treatments. During the 2021 (C) season 60% ET baseline was maintained, but the treatments were changed to 90% ET and 120% ET. Error bars are SEM. N=9. P<0.05; significance was evaluated by two-way analysis of variance.

### Flavonol Content

Flavonol responses broadly mirrored those of anthocyanins, with the 60% ET baseline declining following HW events while supplemental treatments maintained or increased content. However, unlike anthocyanins, flavonol differences between treatments were largely absent at commercial harvest in all three years.

In 2019, the 60% ET baseline declined in flavonol content following HW2 while both supplemental treatments continued to increase through harvest (Figure 3A). Despite mid-season separation, no significant difference was observed between the 60% and 180% ET treatments at commercial harvest. In 2020, treatment differences emerged at the second sampling point in the absence of a HW, then widened during HW3, when the 60% ET baseline fell significantly below the 120% ET treatment (Figure 3B). By HW4 the 120% ET had significantly higher flavonol content than both other treatments, yet no significant differences remained at commercial harvest. In 2021, the 60% ET baseline had the lowest flavonol content at the second sampling point (August 5th) but had surpassed the supplemental treatments by August 17th (Figure 3C). Significant differences between the 60% and 90% ET treatments persisted through September 8th but were absent at commercial harvest.

**Figure 3.**
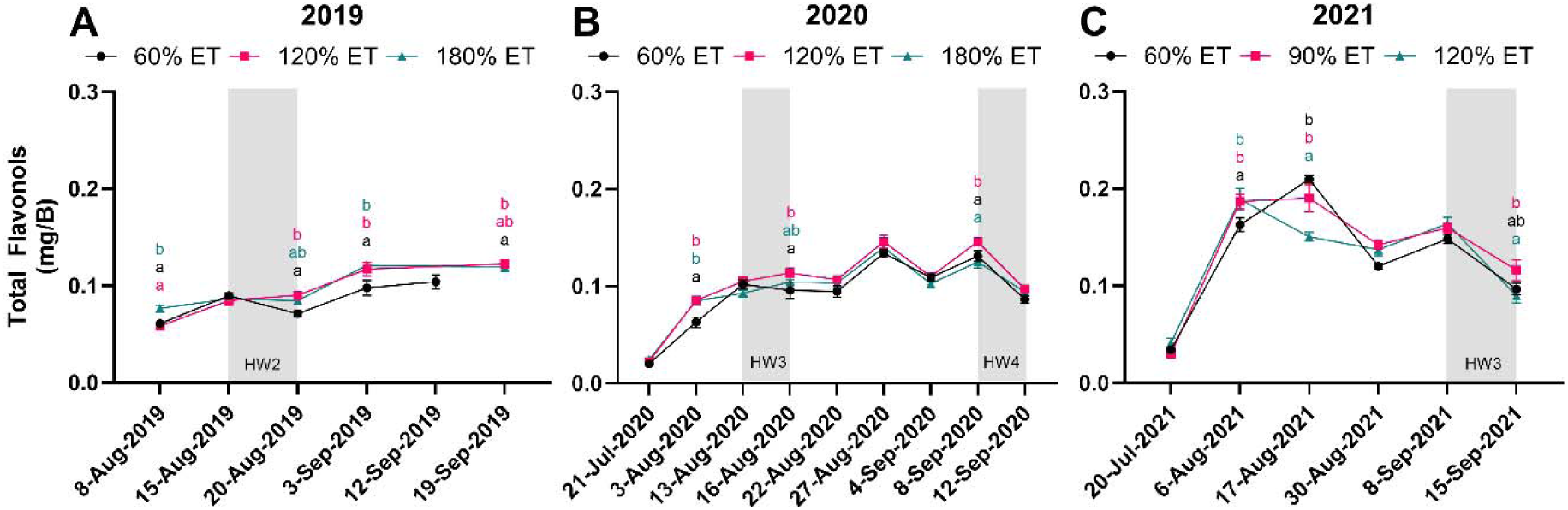
Total flavonol content (milligrams of quercetin-3-glucoside equivalents per berry) throughout ripening during the 2019 (A) and 2020 (B) seasons for the 60% ET baseline, 120% ET, and 180% ET treatments. During the 2021 (C) season 60% ET baseline was maintained, but the treatments were changed to 90% ET and 120% ET. Error bars are SEM. N=9. P<0.05; significance was evaluated by two-way analysis of variance.

### Proanthocyanidins Content

Supplemental irrigation generally maintained higher PA levels than the 60% ET baseline, with HW events driving marked declines across all treatments. Results were less consistent than for anthocyanins and flavonols, with the pattern of treatment separation varying by year.

In 2019, PAs were significantly lower in the 60% ET baseline across all sampling time points, and all treatments declined following HW2 (Figure 4A). In 2020, pre-veraison sampling was included to capture PA synthesis dynamics (Figure 4B). No significant differences were present at the outset, but the 120% ET treatment had significantly higher PAs than the 60% ET baseline following HW2. All treatments then declined markedly at HW3, eliminating differences through commercial harvest, at which point the 180% ET — but not the 120% ET — differed significantly from the 60% ET baseline. In 2021, the 90% ET treatment had significantly more PA material than both other treatments at the start of sampling, but the 120% ET had the highest PAs by the end of HW2 (Figure 4C). Few differences were observed between treatments from August 6th until commercial harvest, when a decline in the 60% ET baseline left the 90% ET treatment with significantly higher PAs.

**Figure 4.**
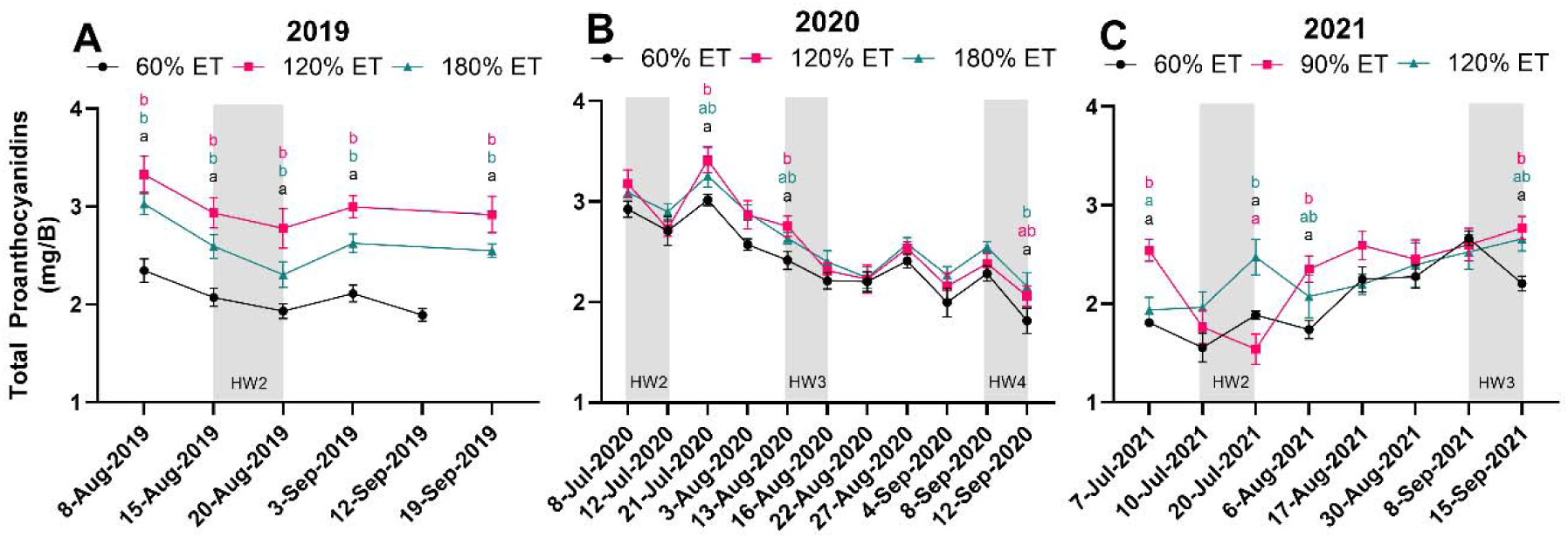
Total proanthocyanidin content (milligrams of epicatechin equivalents per berry) throughout ripening during the 2019 (A) and 2020 (B) seasons for the 60% ET baseline, 120% ET, and 180% ET treatments. During the 2021 (C) season 60% ET baseline was maintained, but the treatments were changed to 90% ET and 120% ET. Error bars are SEM. N=9. P<0.05; significance was evaluated by two-way analysis of variance.

## Discussion

Total soluble solids, acidity, and flavonoids were all significantly impacted by irrigation amount during heatwaves, but the direction and magnitude of those effects depended on whether the HW occurred before or after veraison. This interaction between timing and amount is a recurring theme across all measured parameters and is discussed for each compound class below.

In the case of TSS, pre-veraison HWs enhanced sugar accumulation in the 60% ET baseline, consistent with increased phloem translocation under source-limiting conditions. This was observed in 2020 and 2021; samples were not taken pre-veraison in 2019. A faster rate of TSS accumulation in the 60% ET baseline was also observed when HWs took place post-veraison. Galeano et al. documented a significant decline in stomatal conductance and assimilation rates in the 60% ET baseline during HWs, suggesting reduced carbon availability for sugar loading. ^57^ However, the significantly lower yield and berry weight in that treatment could alternatively explain the accelerated TSS accumulation via concentration effects rather than enhanced loading per se, accounting for the harvest-period differences observed in 2019 and 2020.

Pre-veraison heatwaves impacts on titratable acidity were related directly to the amount of irrigation applied. Although pH differences between treatments were negligible, titratable acidity increased following pre-veraison HWs in the 60% ET baseline, consistent with enhanced (L)-malate synthesis, which is upregulated during this developmental phase and accelerated under warmer conditions.^12^ Post-veraison HWs produced the opposite response in all years: pH increased and titratable acidity declined in the 60% ET baseline, consistent with accelerated malate degradation via elevated respiration rates.^7,14^ Supplemental irrigation dampened both of these responses, with higher irrigation amounts producing progressively smaller deviations from pre-HW titratable acidity values, suggesting that maintaining vine water status buffers the organic acid pool against HW-driven perturbations regardless of developmental timing.

Berry weights are sensitive to irrigation amount during cell differentiation prior to veraison ^6^, and supplemental water was applied during HWs that occurred in the pre-veraison window across all three years (Figure 1). The 120% ET treatment (2019 and 2020) and 90% ET treatment (2021) produced the highest yields, though these did not differ significantly from the 180% ET treatment (Table 2). Because both yields and berry weights differed significantly between the 60% ET baseline and supplementally irrigated treatments, the confounding of flavonoid results is possible. However, lower yields in red winegrapes are generally associated with increased flavonoid contents, and expressing results on a per berry rather than per volume basis accounts for dilution effects from larger berries; the influence of crop load on the present flavonoid results is therefore considered minor.

Overall, TSS, titratable acidity and yield showed a consistent negative response to lower water availability during HWs, and the 180% ET treatment provided no meaningful additional benefit over moderate supplemental irrigation, a pattern that recurs across flavonoid compound classes.

### Irrigation amount, timing, and anthocyanin retention

The interaction between irrigation amount and HW timing was most clearly expressed in anthocyanin responses. The mechanistic basis for the protective effect of supplemental irrigation operates through three interacting pathways, each of which is modulated by developmental stage.

Supplemental irrigation may support anthocyanin accumulation during heat waves through several mechanisms. First, maintaining vine water potential sustains evapotranspiration, reducing canopy and cluster temperatures — well-watered vines have been shown to maintain canopy temperatures approximately 7–8°C lower than water-stressed vines during heat events. ^51,58^ By keeping cluster temperatures closer to the biosynthetic optimum (∼25–30°C), irrigation suppresses heat-driven pigment degradation, an effect most critical post-veraison when the berry is most susceptible to heat-driven pigment loss. Second, moderate water deficit promotes ABA accumulation, which upregulates anthocyanin biosynthesis; however, under the combined heat and water stress of deficit irrigation during heat waves, this signal is overwhelmed and catabolism exceeds biosynthesis. ^59^ Supplemental irrigation maintains vines within the hydraulic and thermal range where ABA signaling supports net accumulation. The absence of additional benefit at high irrigation rates is consistent with stomatal limitation becoming rate-limiting for thermal buffering once evapotranspiration is maximized, and with excess water supply attenuating the mild deficit stimulus that promotes biosynthesis. Third, while phloem dominates berry water supply after veraison, xylem backflow continues and irrigation status retains measurable influence over berry water potential throughout ripening ^60,61^, remains physiologically relevant to anthocyanin metabolism even late in the season.

The interaction of these mechanisms with HW timing explains the treatment × timing patterns observed across all three seasons. During all pre-veraison and early post-veraison HWs, the 60% ET baseline exhibited anthocyanin declines consistent with degradation, while supplementally irrigated treatments showed little or no loss, with 120% and 180% ET performing similarly. This confirms that a moderate irrigation amount is sufficient to engage the thermal buffering and ABA-signaling mechanisms described above; excess water at 180% ET provides no further protection. Pre-veraison HWs may also exert lasting effects on post-veraison anthocyanin accumulation: the 2019 season, which had only one HW before veraison, reached the lowest peak anthocyanin content of the three seasons, while 2020 and 2021, each having multiple pre-veraison HWs, achieved higher peak contents across all treatments. This is consistent with prior work showing that pre-veraison heat stress can modulate post-veraison anthocyanin capacity ^34^, and suggests that HW frequency before veraison may influence the upper bound of achievable anthocyanin content — a relationship warranting further controlled study.

The picture is more complex for late-season HWs. In 2020 and 2021, HWs near commercial harvest led to anthocyanin degradation across all treatments, and in 2021 an apparent reversal emerged in which irrigated treatments showed greater proportional anthocyanin loss during the final late-season HW. Two factors account for this. First, by the time of late-season HWs, anthocyanin biosynthesis has largely ceased, ^33,35^ leaving degradation as the dominant process in all treatments. The larger anthocyanin pool preserved by supplemental irrigation through earlier events therefore presents more substrate for thermally driven peroxidase catabolism, producing greater proportional losses during the final HW even as irrigated treatments retained more anthocyanin at harvest in absolute terms. Second, in 2021 emergency irrigation was applied to all treatments during a severe mid-season event, temporarily equalizing vine water status and diminishing the thermal buffering advantage of supplemental irrigation during the subsequent late-season HW. Additionally, 14 days with temperatures at or above 35°C in 2021 (Figure 1) — none of which met the methodological HW criterion — likely produced cumulative degradative effects that reduced treatment differentiation near harvest.^32,33,62^ Together, these results indicate that irrigation amount has the greatest influence on anthocyanin preservation during early and mid-ripening HWs, and that its effectiveness is attenuated for late-season events when biosynthesis is no longer active.

### Irrigation amount, timing, and flavonol retention

Flavonols share the heat-sensitivity of anthocyanins but differ in their biosynthetic regulation, which shapes how irrigation influences their accumulation. Unlike anthocyanins, flavonol synthesis is driven primarily by light exposure throughout canopy development rather than being concentrated in a post-veraison biosynthetic window, and ABA plays a less direct regulatory role. ^37,63,64^ As a result, heat-driven degradation at any stage tends to produce non-recoverable losses — most evident in 2019, where flavonol content never recovered following the second heat wave. This pattern distinguishes flavonol dynamics from those of anthocyanins, which can partially recover between events if biosynthesis remains active. Given the importance of flavonols to color stability in young wines through copigmentation with anthocyanins, their retention in the berry is of direct relevance to fruit and wine quality.

The consequences of this distinction are evident in the irrigation treatment responses. The 60% ET baseline consistently showed the lowest flavonol contents across all three seasons, despite the likelihood of greater fruit light exposure in that treatment due to reduced vegetative growth — supporting the interpretation that thermal degradation rather than differences in light-driven synthesis drove treatment differences, with higher canopy temperatures in deficit-irrigated vines promoting greater heat-driven catabolism. ^33,57^ Supplemental irrigation at 120% and 180% ET effectively limited flavonol degradation during heat waves occurring in the middle of the ripening phase, with no significant difference between those two treatments — consistent with a stomatal-limitation ceiling on thermal buffering. However, for heat waves at or near commercial harvest, flavonol degradation occurred across all treatments, and treatment differences at harvest were absent in two of three seasons — with the sole exception of 2019, where irrigated treatments retained significantly more flavonols, though again without difference between the 120% and 180% ET levels. This convergence differs from the anthocyanin outcome, where irrigated treatments retained significantly more pigment at harvest in most seasons, and may reflect either the inherently greater stability of flavonol glycosides compared to anthocyanins, or cumulative erosion of flavonol stocks across all treatments by repeated late-season heat exposure. Either way, the protective window in which irrigation amount matters for flavonols is narrower than for anthocyanins, largely confined to mid-ripening heat waves, with neither moderate nor high irrigation providing meaningful flavonol protection — and by extension, copigmentation capacity — during late-season events.

### Irrigation amount, timing, and proanthocyanidin extractability

The synthesis of PAs has been shown to occur between flowering and the onset of veraison.^22^ In previous experiments, light incidence on the cluster has been shown to impact their content, while temperature fluctuation and diurnal changes appear to have less of an effect.^22,48,65^ Furthermore, experiments focusing on pre-veraison deficit irrigation have suggested that the synthesis and content of skin-derived PAs is not strongly dictated by water status during that period. Due to this, the synthesis of PAs is distinctly different from that of anthocyanins, as it has been well documented that deficit irrigation practices lead to increased anthocyanin contents. One mechanism may be through increased ABA production, which has been shown to have a significant effect on anthocyanins when applied exogenously.^66,67^ In this study, heat waves occurred during the flowering-to-veraison window in all three seasons, meaning that PA synthesis was ongoing during some heat events. This is particularly relevant in 2021, where pre-veraison heat waves coincided with active synthesis, and may explain the anomalous pattern observed in that season.

Previous work has shown a decrease in PA material across the course of the growing season in exhaustive extracts, where there was a higher PA content at the beginning of sampling than at commercial harvest; this was true for the 2019 and 2020 vintages. However, there was a slight increase in PAs in 2021. It is possible that there were vintage-to-vintage differences in polysaccharides, or cell wall material, leading to impacts on the extractability of these compounds, but this experiment was not designed to capture results regarding this. ^68^Responses to varying irrigation during heat waves showed broad consistency across vintages, though with some seasonal variation. At commercial harvest, the baseline treatment consistently had the lowest overall PA contents, implying that deficit irrigation combined with heat stress may impact extractable PA contents during the harvest window even after the period of active synthesis. In 2019 and 2021 there were no significant differences in PA content between the higher irrigation treatments at harvest, consistent with a stomatal-limitation ceiling on any protective effect of supplemental water. However, in 2020 the 180% ET treatment had significantly higher PAs than the 60% ET baseline at harvest, while the 120% ET treatment did not differ significantly from either, suggesting that the dose-response relationship between irrigation amount and PA retention was not consistent across seasons. This suggests that the application of water in excess is not universally beneficial for the mitigation of PA losses, which has implications for efforts to reduce water use under continued pressure on these resources.

A growing number of publications have shown the impacts of heat stress and water scarcity on the synthesis and stability of anthocyanins and flavonols in the skins of wine grapes. In this work, the goal was to understand the impacts of a mitigation technique, already used widely in the industry, to evaluate its effect on flavonoid synthesis and preservation. The results show a clear impact of water application on the loss of polyphenols, especially anthocyanins, during the growing season. However, the data also show that irrigation in excess prior to a late-season heat wave has negligible additional benefits regarding the prevention of polyphenol degradation. Therefore, the results provide an initial roadmap for viticulturists to mitigate the damages of heat waves at key points during the ripening phase, while conserving water. Future efforts should focus on how these results may be impacted under different climatic, soil, and variety-dependent conditions.

## Abbreviations Used

ABA: Abscisic Acid
CWM: Cell wall material
DAD: Diode Array Detector
ET: Evapotranspiration
HW: Heatwave
MDI: Moderate Deficit Irrigation
MCP: Methyl Cellulose Precipitation
PA: Proanthocyanidin
TA: Titratable Acidity
TFA: Trifluoroacetic Acid
TSS: Total Soluble Solids
VRDI: Variable Rate Drip Irrigation

## Acknowledgements

Support from E. & J. Gallo and their staff was critical to the success of the field trial, as well as seed funding from USDA California Climate Hub, CDFA Specialty Crop Block Grant Award No. 19-0001-013-SF, the American Vineyard Foundation, the Department of Viticulture & Enology and the College of Agricultural and Environmental Sciences at UC Davis.

