## Supplemental Materials for "Enhanced Irrigation during Extreme Heat Events Preserves Anthocyanins in Cabernet Sauvignon"

**Supplementary Data.**

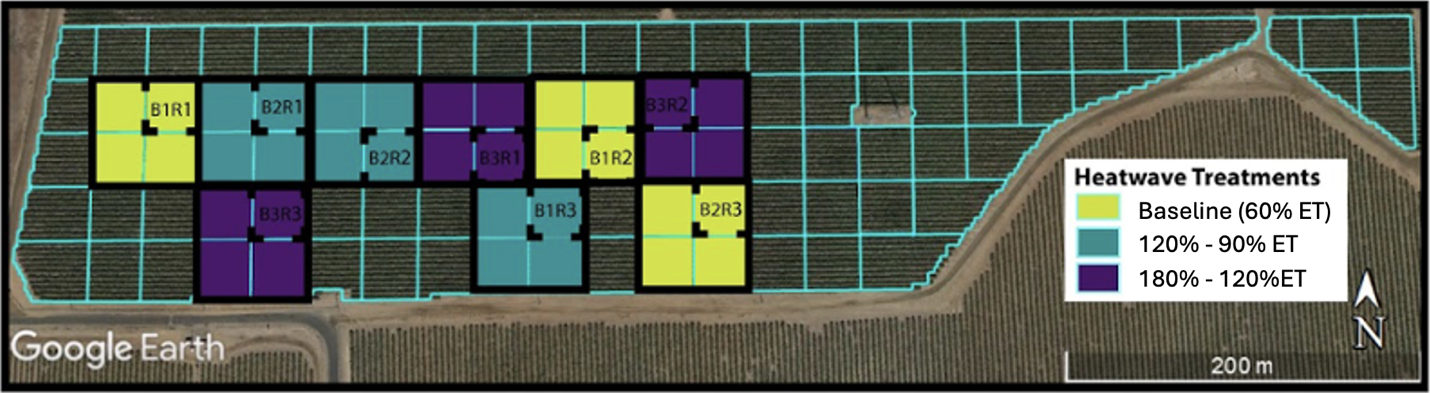
**Figure S1. Example of site & treatments.**  Layout of the vineyard study site and experimental design of irrigation treatments, with the 30 x 30 meter squared 2019 plots indicated with a dashed line within the 60 x 60 meter squared 2020 and 2021 plots.

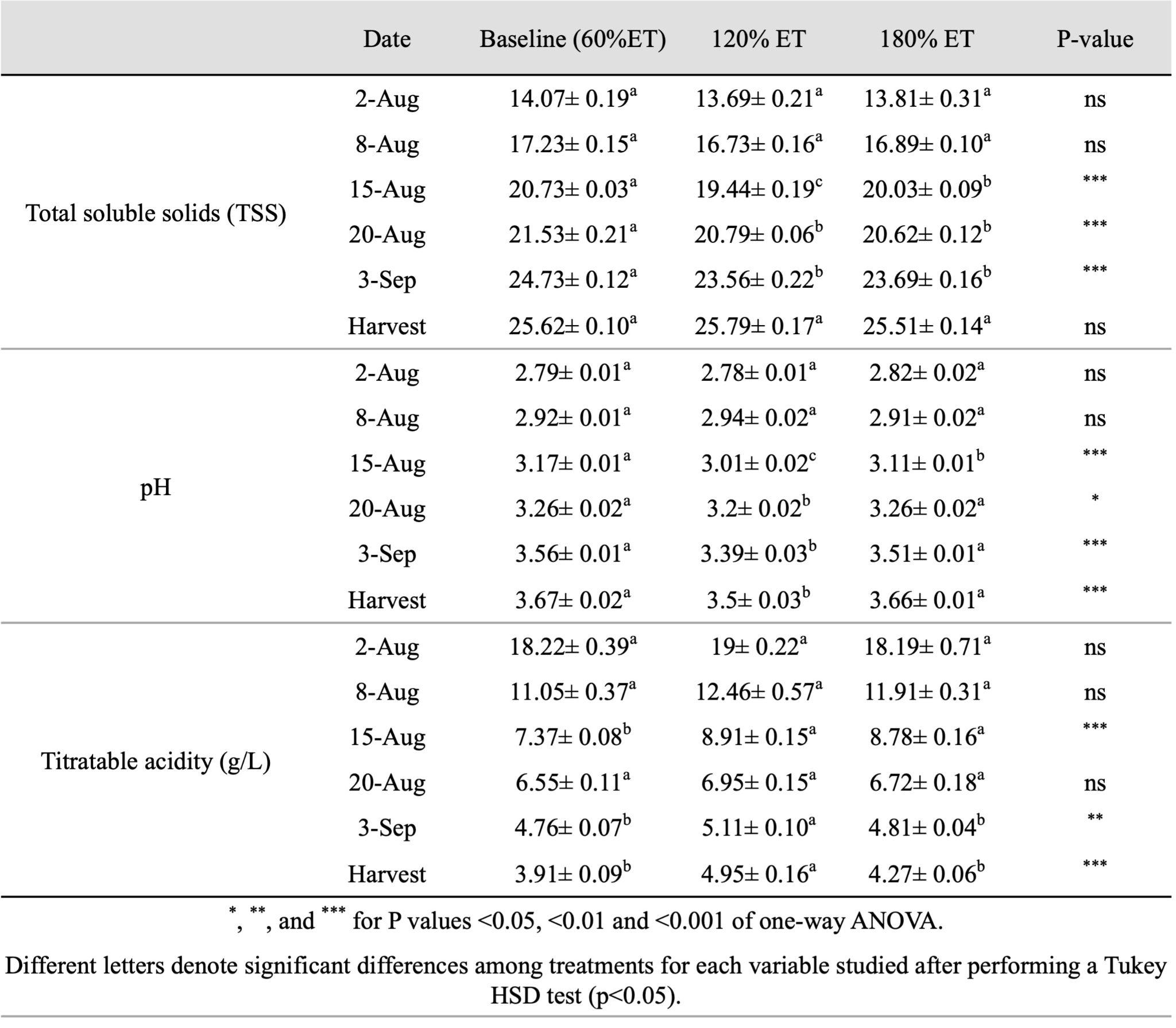
**Table S1. Complete 2019 Primary Chemistry Table.** Dynamics of berry primary metabolites during ripening in the 2019 season of treatments exposed to differential irrigation (Baseline (60% ET), 120% ET, and 180% ET). Total soluble solids expressed as Brix, pH, and titratable acidity expressed in g/L of tartaric acid equivalents.

**Table S2. Complete 2020 Primary Chemistry Table.** Dynamics of berry primary metabolites during ripening in the 2020 season of treatments exposed to differential irrigation (Baseline (60% ET), 120% ET, and 180% ET). Total soluble solids expressed as Brix, pH, and titratable acidity expressed in g/L of tartaric acid equivalents.

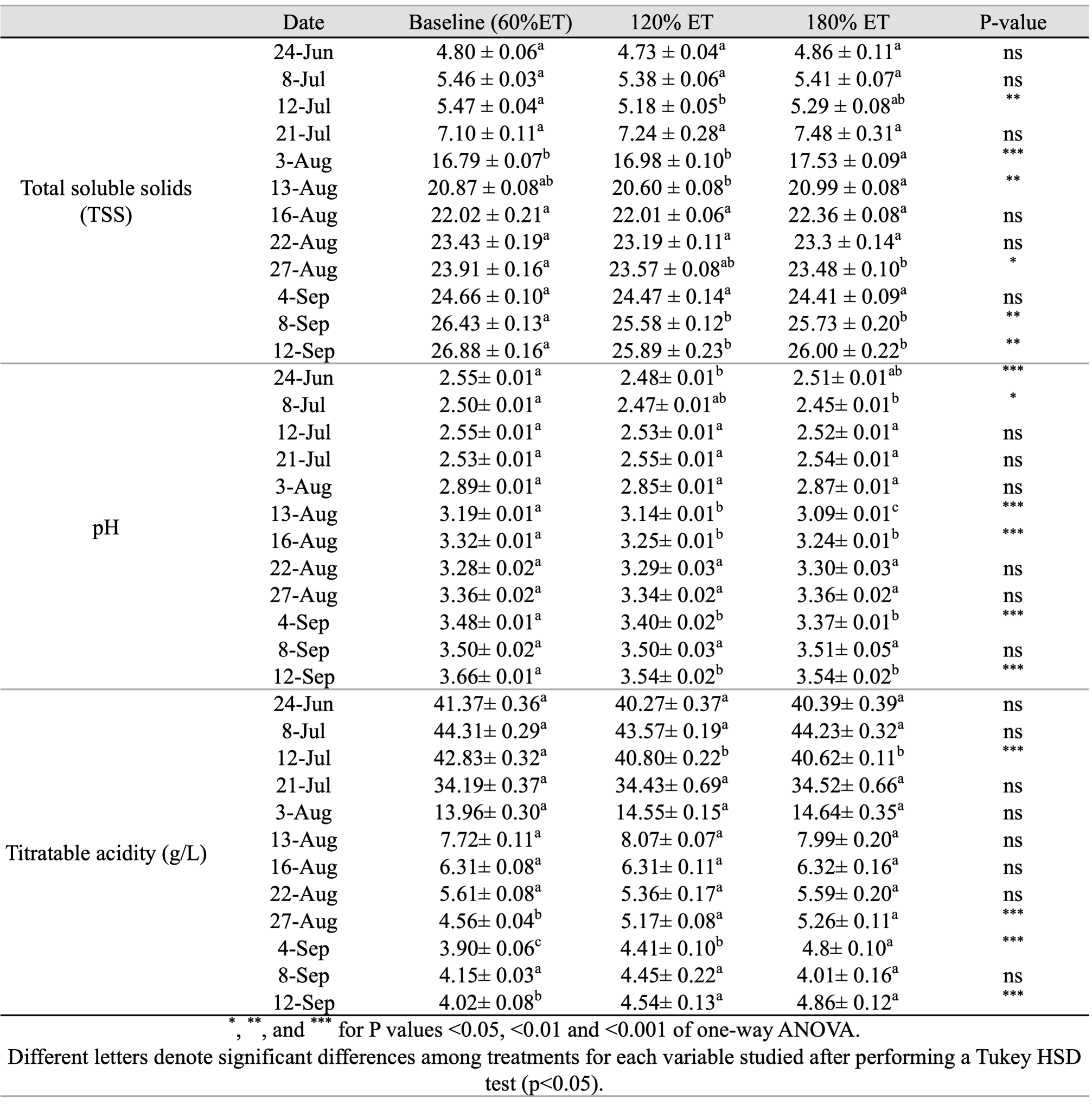

**Table S3. Complete 2021 Primary Chemistry Table.** Dynamics of berry primary metabolites during ripening in the 2021 season of treatments exposed to differential irrigation (Baseline (60% ET), 90% ET, and 120% ET). Total soluble solids expressed as Brix, pH, and titratable acidity expressed in g/L of tartaric acid equivalents.

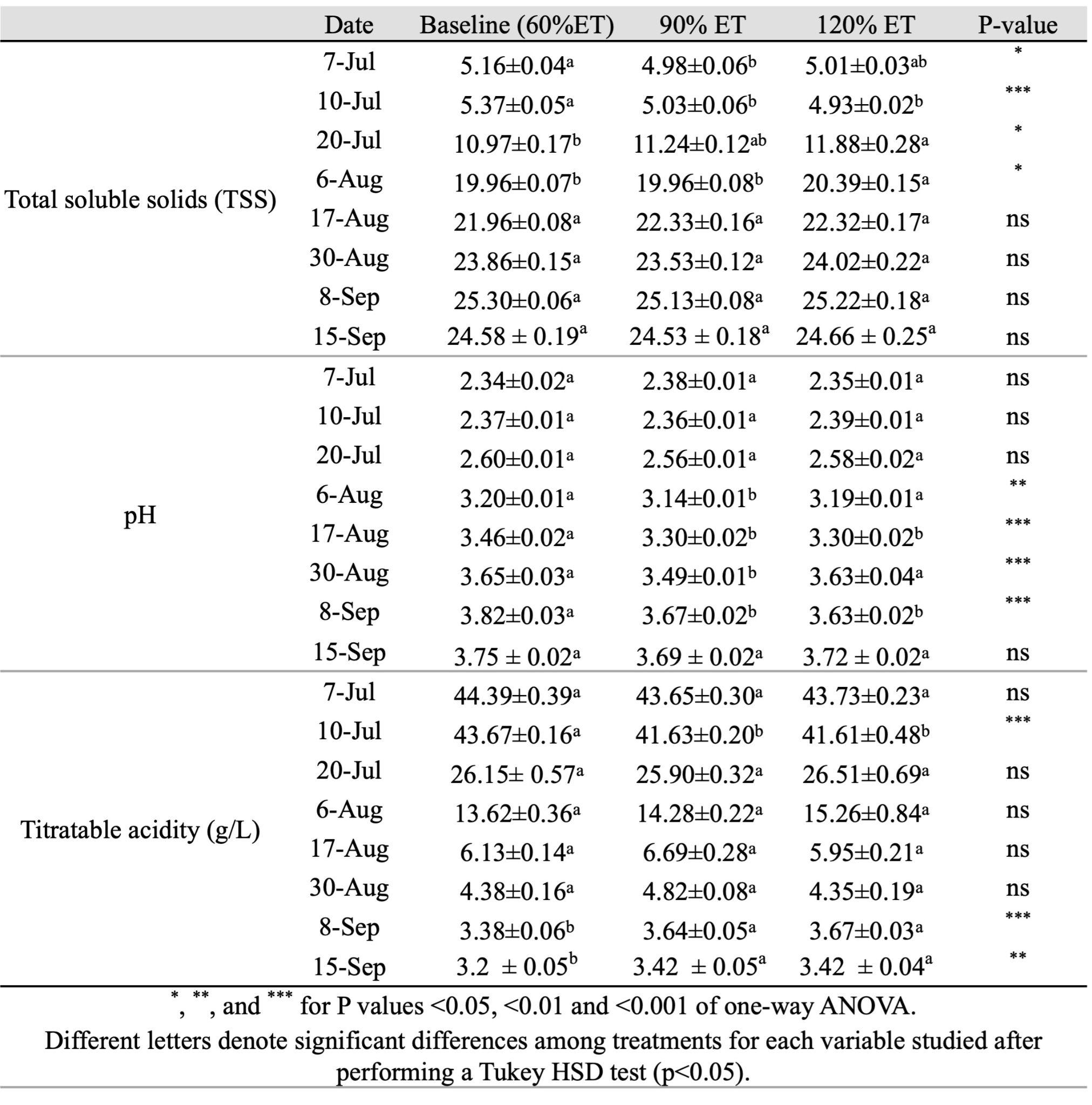
